# Modeling the Shape of Synaptic Spines by their Actin Dynamics

**DOI:** 10.1101/817932

**Authors:** Mayte Bonilla-Quintana, Florentin Wörgötter, Christian Tetzlaff, Michael Fauth

## Abstract

Dendritic spines are the morphological basis of excitatory synapses in the cortex and their size and shape correlates with functional synaptic properties. Recent experiments show that spines exhibit large shape fluctuations that are not related to activity-dependent plasticity but nonetheless might influence memory storage at their synapses. To investigate the determinants of such spontaneous fluctuations, we propose a mathematical model for the dynamics of the spine shape and analyze it in 2D — related to experimental microscopic imagery — and in 3D. We show that the spine shape is governed by a local imbalance between membrane tension and the expansive force from actin bundles that originates from discrete actin polymerization foci. Experiments have shown that only few such polymerization foci co-exist at any time in a spine, each having limited life time. The model shows that the momentarily existing set of such foci pushes the membrane along certain directions until foci are replaced and other directions may now be affected. We explore these relations in depth and use our model to predict shape and temporal characteristics of spines from the different biophysical parameters involved in actin polymerization. Reducing the model further to a single recursive equation we finally demonstrate that the temporal evolution of the number of active foci is sufficient to predict the size of the model-spines. Thus, our model provides the first platform to study the relation between molecular and morphological properties of the spine with a high degree of biophysical detail.

**Author summary:** Synaptic spines are post-synaptic contact points for pre-synaptic signals in many cortical neurons and it has been shown that synaptic transmission is correlated with spine size. However, spine size and shape can vary quite strongly on short time scales and it is currently unknown how these shape variations emerge. In this study we present a biophysical model that links spine shape fluctuations to the dynamics of the spine’s actin-based cytoskeleton. We show that shape fluctuations arise from the fact that fast actin polymerization in a spine is a discrete process happening at only few polymerization foci. Life and death of these foci determine from moment to moment how the membrane bulges or retracts. We provide an in-depth analysis of this effect for a large set of biophysical parameters and quantify the spatial-temporal structure of the spines. Our model, thus, provides a quantitative characterization of the link between spine morphology and the underlying molecular processes, which forms an essential step towards a better understanding of synaptic transmission during steady state but also during synaptic plasticity.

## Introduction

Dendritic spines are small protrusions from neural dendrites, which form the postsynaptic part of most excitatory synapses in the cortex [1]. One of the central paradigms of neuroscience is that synapses store memories by changing their transmission efficacies during learning [2] and it has been shown that magnitude as well as synaptic transmission efficacy correlate with size and shape of the spines. This has been mostly studied using the volume of the spine head ([3–6], more details in [7]) providing evidence for a link between spine-morphological and synaptic-functional properties. However, it recently became clear that most of the dynamic properties of changing spine volumes emerge from spontaneous spine specific processes that are not determined by the activity of the pre- or postsynaptic neuron [8–10]. As such spontaneous fluctuations could affect memory functions due to the above described link [11], a thorough understanding of their characteristics and underlying processes is necessary. Experiments imaging the shape of dendritic spines can provide snapshots at distinct time points, but mathematical models are needed to bridge between these time points and to understand shape fluctuations and their properties. However, so far only phenomenological models have been proposed [9, 12–14] that describe fluctuation coarsely on a timescale of days. Here, we take a different approach by modelling the fast actin dynamics underlying shape fluctuations. This approach also allows us to explore the influence of the molecular and mechanical processes involved and to make predictions on the fluctuations when their properties vary.

The spine shape is determined by its cytoskeleton, the main component of which is actin. Actin is a globular protein (G-actin), which can assemble into filamentous polymers (F-actin). These polymers undergo a continuous treadmilling process (Fig. 1A; see, e.g., [15–17] for details): G-actin with bound ATP is added preferentially to the barbed (+) ends of the filament (see for example added monomer marked with P in Fig. 1A), while at the pointed (-) end older actin monomers of the filament are mostly depolymerized. Thus, actin filaments are polar structures with one end growing more rapidly than the other. This asymmetry between barbed and pointed end is further strengthened when the ATP bound at actin filaments hydrolyzes to ADP, which promotes disassembling of the pointed ends by severing proteins, such as cofilin (D in Fig. 1A), when the pointed end is in an uncapped state (U in Fig. 1A). Following this, the disassembled cofilin-ADP-actin dissociates to cofilin and an ADP-actin monomere and finally, profilin catalyzes the exchange of ADP to ATP and the resulting ATP-actin is again available for the polymerization process at the barbed end (omitted in Fig. 1A). Additional to this treadmilling process, complexes such as ARP2/3 can induce branching of a filament whose two daughter-filaments have uncapped barbed ends (B in Fig. 1A) and capped minus ends. Moreover, barbed ends can become unable to polymerize G-actin due to capping proteins (C in Fig. 1A).

**Fig 1.**
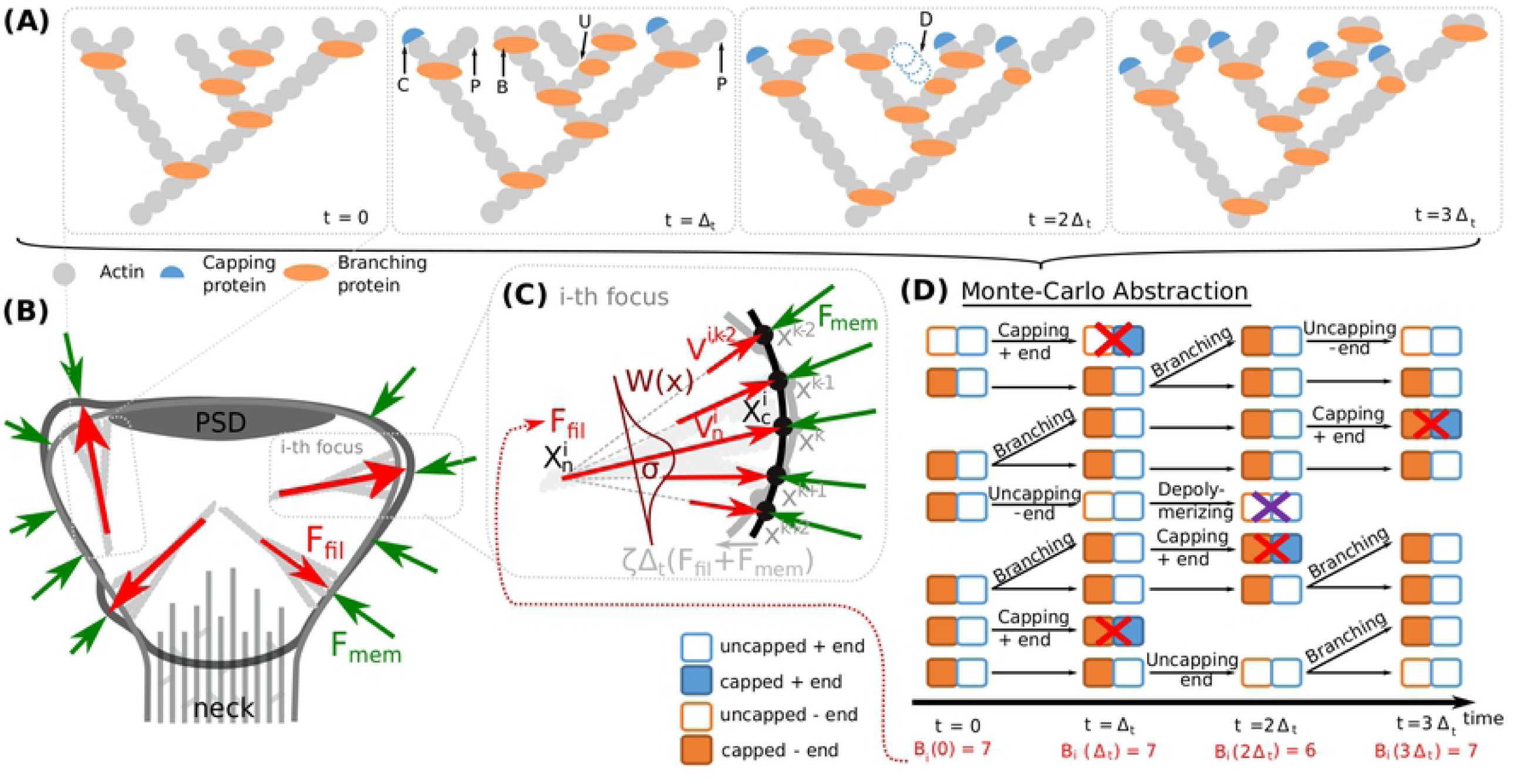
Components of the proposed spine fluctuation model. (A) Schematic picture of an actin filament at successive time-points. Additional to polymerization of new actin monomers (P) at the barbed ends, other events can occur in actin filaments, such as: branching the barbed ends by inserting branching protein ARP2/3 (B), capping barbed ends with capping proteins (C), uncapping minus end (U) and depolymerizing uncapped minus ends (D). (B) Our model for spine fluctuation assumes that the shape of the membrane is determined by the membrane forces **F**_*mem*_ resisting bending and stretching and the forces generated by actin polymerization **F**_*fil*_ at a few foci. (C) Actin filaments at the foci are considered to extent laterally to the membrane. Hence, force is proportional to the number of barbed ends at the focus and attenuated by a spatial kernel *W* (*x*). The membrane is simulated by a discrete mesh (here depicted by dots) that move every time step proportional to imbalance of the acting forces (black membrane grey → membrane). (D) The dynamics of actin in a focus are abstracted to a Monte Carlo model describing the state of the barbed and pointed ends of any filament. We depict these state representations for the time course shown in A. During simulations, the transitions between different states happen according to the processes described in A with defined rates. Details see main text.

**Fig 2.**
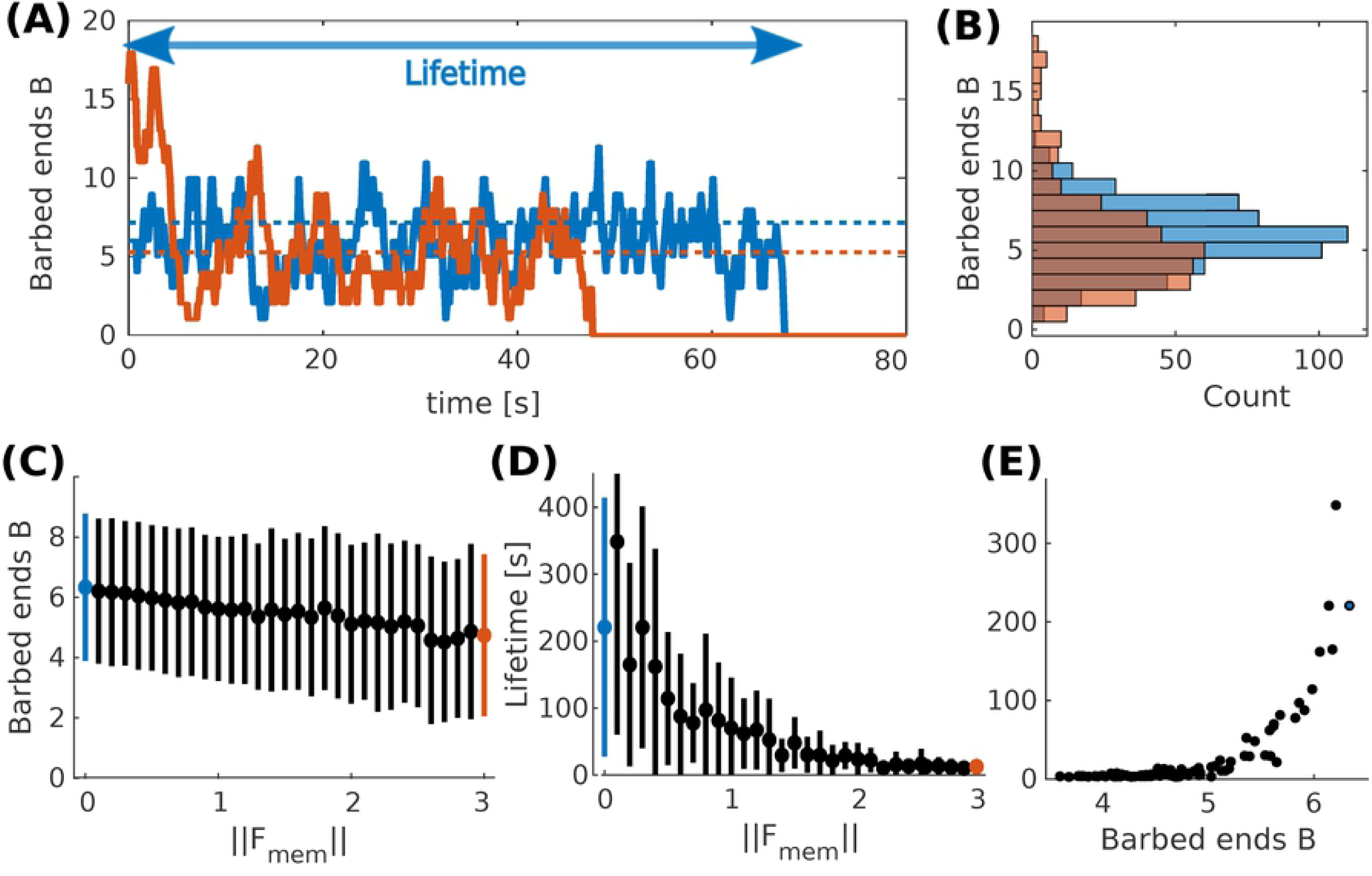
Actin polymerization focus. (A) Evolution of barbed ends over time at a single actin polymerization focus. Blue solid line correspond to ‖**F**_*mem*_‖ = 0pN and orange solid line to ‖**F**_*mem*_ ‖= 3pN. Dotted lines indicate theoretical mean values.. Arrow indicates the lifetime of the focus corresponding to the blue curve. (B) Occurrence frequency (x-axis) of each number of barbed ends (y-axis) over all time-steps (of length Δ_*t*_) in (A). Colors as in (A). (C) Mean and standard deviations of mean number of barbed ends *B* over 50 simulations each for different values of ‖**F**_*mem*_‖. (D) Same for focus lifetimes. Blue and red dots mark the force values used in panel (A) and (B). (E) Mean lifetime over mean number of barbed ends over 50 simulations for varying force values.

Although the treadmilling process in dendritic spines occurs at different velocities, two distinctive pools of F-actin can be identified [18]: The static pool, which has a slow treadmilling velocity and is localized at the base of the spine head whilst the dynamic pool treadmills faster and is found at the tip of the spine head. Honkura et al. [18] suggest that these pools have different functions: the static pool gives stability to the base of the spine, while the dynamic pool causes spine expansion due to the higher rate of actin polymerization resulting from the fast treadmilling velocity. Such fast treadmilling, however, is not occurring uniformly distributed over the whole spine, but at discrete foci of actin polymerization [19]. There are usually only a handful of those foci in one spine, which are well-separated from each other and can be identified from their increased polymerization rate. It can be assumed that these foci generate the main expansive force that underlies shape fluctuations, which are usually inhomogeneous and asymmetric.

Mathematical models that link actin activity to such asymmetric spine fluctuations are, however, missing so far. Although models of the actin treadmilling process have been derived [16] and adapted to the conditions in the dendritic spine [17], they have not been connected to spine shape. To evaluate how shape is influenced by actin dynamics, one has to consider not only the forces created by the filaments, but also the counteracting forces from the lipid membrane that encloses the spine. Such models for force generation by actin filaments [20] and their interaction with the membrane have been derived and successfully applied to the movement of bacteria, cell motility ([21–23]; for a review see [24]), and to explain dendritic spine maturation [25]. Yet, most of these models describe the dynamics of the cell shape based on density descriptions of the actin filaments or assume a homogeneous distribution of F-actin. However, considering the comparably small numbers of filaments within the spine (compare [26]) a density description is not applicable. The homogeneity assumption, in turn, entails very regular and symmetric spine shapes, which are not observed in experiment (e.g., [27]) and also not consistent with the existence of actin polymerization foci.

Here, we present a model that considers heterogeneous actin dynamics caused by foci of actin polymerization. We use the forces generated by their treadmilling activity together with the counteracting forces from the membrane and the membrane-mediated coupling to other foci to derive a model of spine membrane shape fluctuations in 2D and its extension to 3D. We show that the properties of spine fluctuations are strongly influenced by the dynamics of filament assembly constituting the determinants of the force generation by actin. The central finding of this study is that spine shape fluctuations can be fully explained by the effect that the small number of polymerization foci leads to a discretization of the outwards pushing-force direction, while their limited life time determines the temporal properties of these fluctuations. Thus, we can also show that spine area evolution can be predicted by the number of polymerization foci. Thus, this model provides the required biophysically detailed basis for future extension into investigation of spine shape changes induced by synaptic plasticity.

## Materials and methods

### Model

Based on the findings of [19], we assume that the spine shape is determined by a small number of distinct foci of actin polymerization (grey filaments in Fig. 1B), for which the processes of treadmilling, branching and capping of the filaments are modelled individually (see Sec. Actin dynamics at individual foci). As a consequence, each focus can have multiple barbed ends generating forces that push the membrane outward (see Sec. Actin-generated force, red arrows in Fig. 1B and C). These forces concur with the inward directed forces generated by the membrane’s resistance against deformation (see Sec. Membrane force, green arrows in Fig. 1B and C). If these forces are locally unbalanced, the membrane moves giving rise to shape fluctuations (transition from black to grey membrane shape in Fig. 1C). To simulate this interaction of membrane shape and forces, we use discrete timesteps and the finite elements method. In particular, the membrane is represented by a mesh of points (or vertices) for which geometrical properties, forces and movements are calculated (see S2 Appendix,S3 Appendix).

#### Membrane mesh initialization and morphological constraints

As stated above, we represent the membrane enclosing the spine by a mesh of vertices *k* ∈ {1, 2, …*n*_*vertices*_} described by their (2 dimensional) position vectors **x**^*k*^. Upon initialization, a polygonal approximation for a circle with radius *r*_*s*_ and centered at the origin of the x-y plane is created. As in this study we focus on the shape fluctuations of mature spines, we also implement two major morphological constraints:

On the one hand, the spine neck of mature spines typically contains heavily interlinked actin bundles which are rather stable and have a much slower treadmilling velocity than those in the spine tips [28]. Along this line, also the spine neck width is largely stable on the here considered timescale of hours [29]. Therefore, we fix the location of mesh-points at the neck throughout the whole simulation. We establish those fixed mesh points during the mesh initialization by selecting all points **x** = (*x, y*) with *y* ≤ *h*_*neck*_ and fixing them to (*x, h*_*neck*_), here we define *y* = *h*_*neck*_ as the value where *x* = *r*_*neck*_ for *y* < 0.

On the other hand, also the movement of the post-synaptic density (PSD) is constrained as it is heavily interlinked with the presynaptic site. Also, the PSD size on unstimulated spines is conserved over the here considered timescale of hours [30]. Therefore, we also fix the mesh-points (*x, y*) with *y* ≥ *h*_*PSD*_ to (*x, h*_*PSD*_), where *y* = *h*_*PSD*_ is the value where *x* = *r*_*PSD*_ for *y* >0.

#### Actin dynamics at individual foci

The amount of force generated by actin polymerization is assumed to depend mainly on the number of active barbed ends at each polymerization focus. Therefore, we reduce the complicated process of filament assembly, branching and capping at each focus *i* to the dynamics of the filaments that have uncapped barbed ends abstracting the exact geometrical properties of the branched tree of actin filaments (Fig. 1D). Each of these active filaments is characterized by the state of its barbed end (normally uncapped) and the state of its pointed end (normally capped or bound to a ARP2/3 complex). The dynamics that change these states are stochastic and simulated in discrete time steps of length Δ_*t*_. Based on Bennett et al. [17], it is assumed that besides F-actin polymerization, i.e., the addition of new G-actin at the uncapped barbed ends, the following processes can occur at each filament:

- Uncapped barbed ends branch by including an ARP2/3 molecule and give rise a new filament with a probability 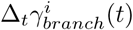.
- Uncapped barbed ends are capped by a capping protein with a probability Δ_*t*_*γ*_*cap*_. Polymerization is not possible when a barbed end is capped such that this barbed end does not generate force. As uncapping occurs very seldom, these filaments are therefore eliminated from the simulation.
- Capped minus ends are uncapped with a probability Δ_*t*_*γ*_*uncap*_.
- Uncapped minus ends are severed with a probability Δ_*t*_*γ*_*sever*_, which leads to the removal of the respective filament.

To simulate this, we iterate through all filaments with uncapped barbed ends within the active actin polymerization foci and the above processes in the indicated order. A process occurs at given filament, when a random number drawn for this filament falls below the indicated probability. Afterwards, the remaining uncapped barbed ends in each polymerization focus *i* are counted and their number *B*^*i*^ is used to calculate the expansive force exerted by that focus. Figure 1D shows an exemplary temporal evolution of the active filaments in one of the polymerization foci, where all of these processes occur. The rate values except *γ*_*branch*_(*t*) (see below) are stated in Table 1.

**Table 1.**
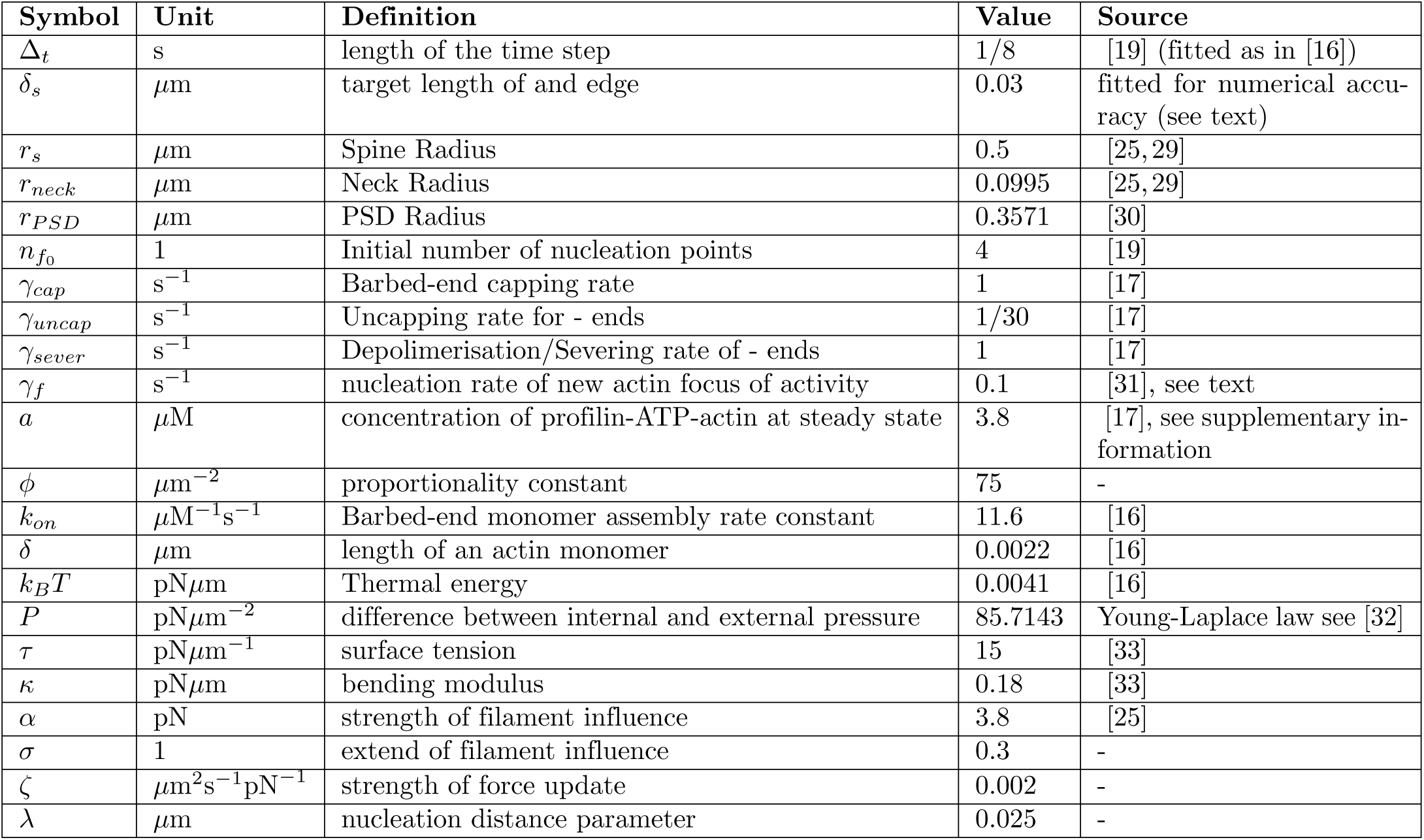
Model Parameter Values.

Following Bennett et al. [17], the branching rate for a filament 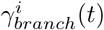 depends on the number of barbed ends *B*^*i*^(*t*) at the respective actin polymerization focus *i* at time *t*:

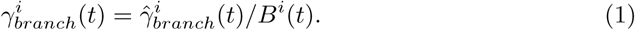

Note, as this is the probability for each single barbed end, the overall branching rate at the focus is 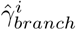. Instead of using a constant 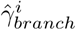 as in [17], we can take changes in the counteracting membrane forces into account. For this, we assume that the branching frequency is proportional to the frequency of adding new monomers. Thus, it becomes proportional to the treadmilling velocity 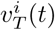, hence, 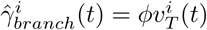. Following Mogilner and Edelstein-Keshet [16], this velocity depends on the number of barbed ends and the membrane force that opposes the outgrowth of the filament:

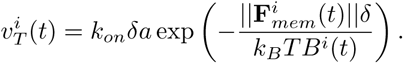

The respective branching rate can be calculated as

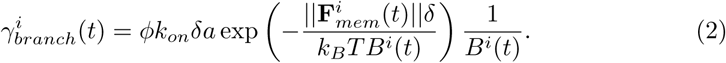

Hereby, *k*_*on*_*δa* is the free polymerization velocity, where *δ* is the length of an actin monomer, *k*_*on*_ the barbed-end monomer assembly rate, and *a* the concentration profilin-ATP-actin available for polymerisation. As we are not modelling plasticity related changes in this study, we can consider *a* as a constant instead of modelling the recycling process explicitly (compare [17], see S1 Appendix).

The free polymerisation velocity is attenuated due to a counteracting membrane force according to the Brownian ratchet theory [20, 34], which takes into account, the absolute temperature *T*, the Boltzmann’s constant *k*_*B*_ and the force 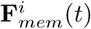 working against polymerization which is generated by the membrane at the *i*th focus center. This membrane-force-dependency of the branching rate generates a feedback between the number of barbed ends and membrane shape.

#### Foci generation and removal

The activity of a focus is determined by its uncapped barbed ends, which can only emerge from other uncapped barbed ends due to branching; hence, foci naturally become inactive and removed as soon as they have no uncapped barbed ends left.

On the other hand, there must also be a mechanism that creates – or nucleates – new foci. In our model, the nucleation of a new focus *i* is implemented in two steps: First, a two dimensional nucleation position denoted by a vector 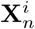 (see Fig. 1C) is selected in the following way: To take into account the assymetrical form of the spine head, we generate a set of 1000 uniformly distributed candidate points inside the spine. From this candidate set, we sort out all points that are not within a distance of 0.1*µ*m from the membrane and that are within 0.1*µ*m from the PSD (based on experimental observations in [19]). Then, one of the remaining *n*_*cand*_ points *j* is selected with probability 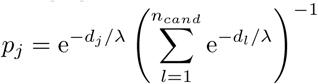, which depends on the distance from the PSD *d*_*j*_ via a scale parameter *λ*. For *λ* → 0^+^ nucleation near to the PSD is favored whereas for *λ* → ∞ the distance to the PSD has no influence.

Second, the primary nucleation direction is randomly selected as the vector pointing from 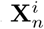 to one of the membrane points that are within 0.1*µ*m. The position of the selected membrane vertex *k* is referred to as the center of the focus 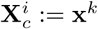. As the foci are relatively short-lived, we assume that this direction does not change over the lifetime of the focus.

#### Actin-generated force

As in Mogilner and Oster [21] we take the propulsive force generated by actin polymerization to be proportional to the number of uncapped barbed ends within each focus. This force is assumed to be acting around the center of an actin polymerization focus, hence, the membrane vertex at position 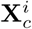. However, as F-actin at the foci is a spatially extended structure, the number of barbed ends *B*^*i*^ at the *i*th focus also affects nearby vertices. We model this by a Gaussian spatial kernel

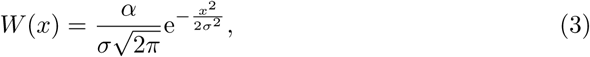

with an amplitude *α* and standard deviation *σ*. The resulting force vector at the vertex *k* (located at **x**^*k*^) is given by

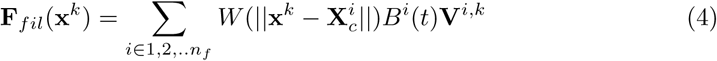

with *n*_*f*_ being the number of currently active actin foci and 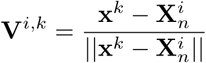 the normalized direction vector of the force from focus *i*.

#### Membrane force

Biological membranes, such as the one confining the spine head, exert forces to resist deformations, especially against being excessively bent or stretched. These forces generated by the membrane (described by a manifold Γ) can be derived from the (Helfrich) free energy [35] given by

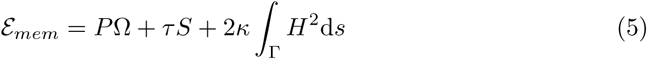

where the membrane’s physical properties are characterized by the difference between internal and external pressure *P*, the line tension (or surface tension in 3D) *τ*, and the bending modulus *κ*. Ω is the area enclosed by the membrane (or volume in 3D), *S* is the boundary length (or surface area) and *H* is the mean curvature. The geometrical properties Ω, *S* and *H* can be derived from the coordinates of the vertices in our discrete mesh describing the membrane shape (see S2 Appendix). The membrane force vector **F**_*mem*_(**x**^*k*^) at vertex *k* is given by

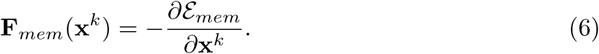

On our discrete mesh, the geometrical properties Ω, *S* and *H* and hence the resulting force can be approximated for each vertex by taking its next neighbors in the mesh into account (see S2 Appendix). Note, however, the approximations of the geometrical properties are only valid when the mesh is dense enough. Therefore, if the vertices move too far apart from each other, our mesh has to be refined (remeshing, see below).

Together with the fixed spine neck and PSD vertices, these forces of the membrane itself give rise to a characteristic “resting shape”, to which the spine converges in the absence of other forces to minimize area, length and curvature (see, e.g., Fig. 4A for rest shapes resulting for different PSD-sizes).

**Fig 3.**
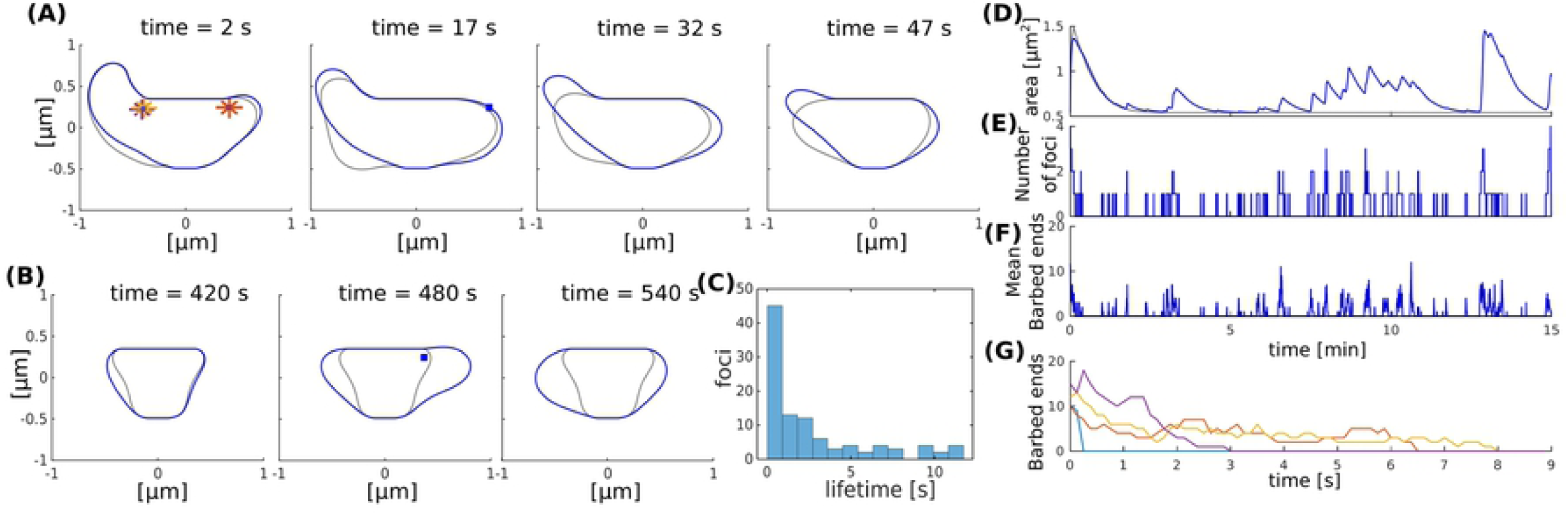
Shape fluctuations of the spine. (A) Snapshots of the spine head shape with (blue line) and without (gray line) nucleation of new actin polymerization focus taken every 15 seconds. Asterisk correspond to the initial nucleation positions **X**_*n*_ and blue squares indicate nucleation positions of the active foci at the indicated time. (B) Same as (A) but taken every 60 seconds. (C) Occurrence frequency of foci lifetime over all time-steps (of length Δ_*t*_) in simulation from panels (D-F). (D-F) Evolution of the spine area (D), number of foci (E) and mean number of barbed ends (F) over time. Gray and blue lines represent the simulation without and with nucleation of new foci respectively. (G) Evolution of the number of barbed ends in each actin focus (color-coded) in the simulation without nucleation of new foci.

**Fig 4.**
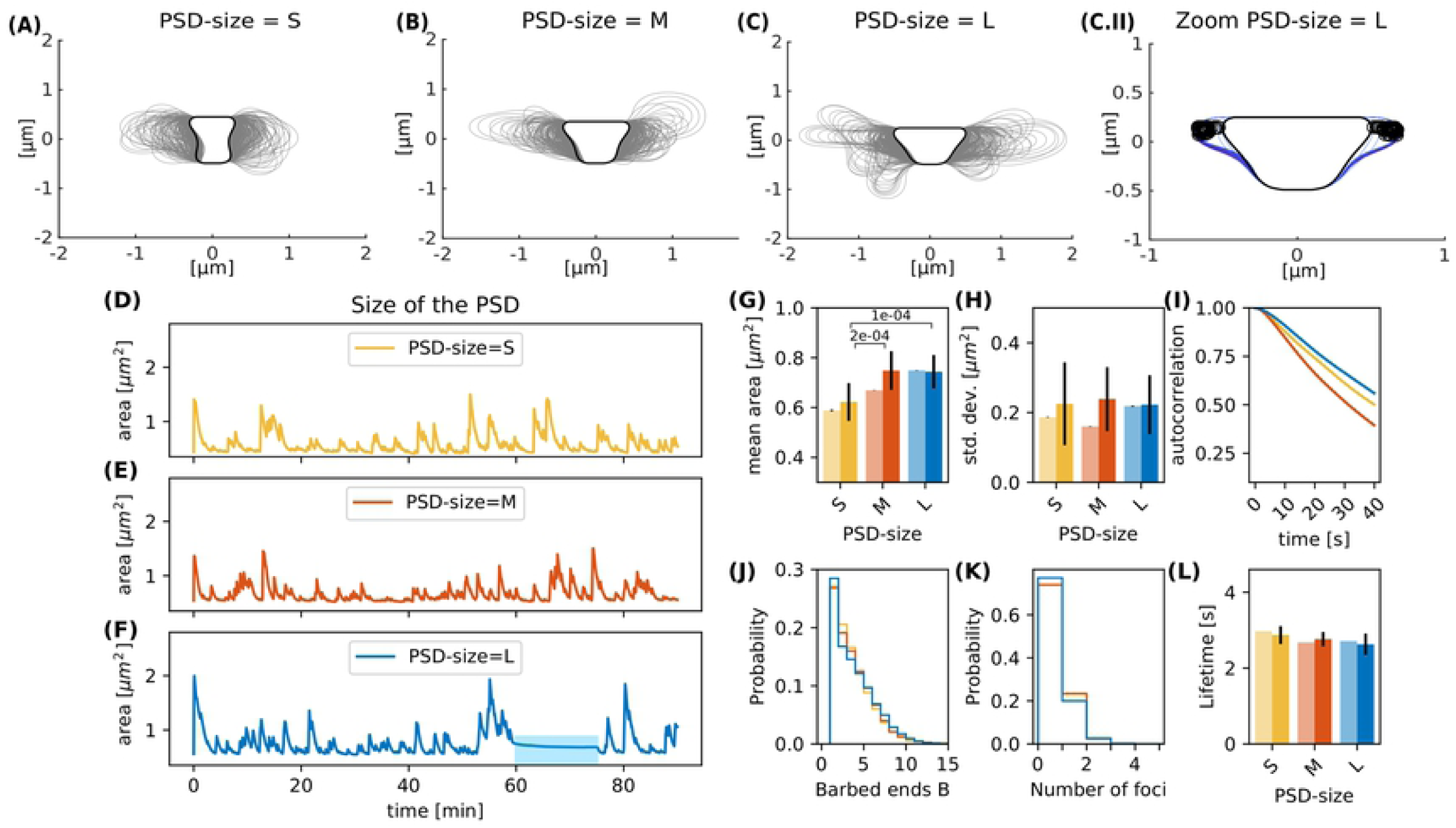
Spines with varying PSD size. Spine shapes (gray lines, plotted every 15 seconds) over 90 minutes of simulation for (A) small, (B) medium and (C) large sized PSDs. Black line corresponds to the resting shape. (C.II) Shapes of the large PSD-size spine between minute 60 and 70 (color-coded from dark to light blue). Black open circles correspond to the nucleation points. (D-F) Evolution of the spine area over time. (G) Temporal mean over area in 90 minute simulation (pale bars, errors are standard deviations over 50 bootstraps) and average of temporal means over fifteen 15 minute simulations (saturated color, errors are standard deviations of the mean) over different PSD sizes. The p-value for significant Welch-tests is indicated. (H) Same as (G) but for standard deviation of the area fluctuations. (I) Autocorrelation functions for the area fluctuations in the 90 minute simulations. (J) Histogram of the mean number of barbed ends at the polymerization foci at each time-step of the 90 minute simulation. (K) Histogram of the active actin polymerization foci. (L) Mean lifetime of a focus, color-coded as in (G).

#### Membrane movement

In the presence of both actin and membrane generated forces, the spine shape is determined by a balance between them. If the forces are unbalanced at one of the mesh vertices, they will generate a movement of that vertex and deformation of the membrane. For simplicity, we here assume that the motion of the vertex *k* is proportional to the net force with a proportionality constant *ζ*. The displacement of vertex *k* in time is given by

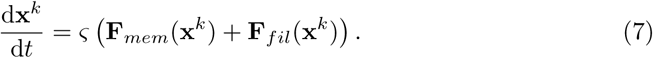

Also this equation is implemented in discrete time-steps. For this we use a classical Runge-Kutta algorithm, in which we keep 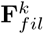 constant during the whole calculation step. However, the interaction between neighboring membrane points can still give rise to diverging oscillations. Therefore, if the membrane displacement in a single time step exceeds a certain displacement tolerance (*d*_*tol*_ = 0.0001*µ*m), we split this time step in two intervals and calculate the displacement of all membrane vertices in each of them until the displacement is smaller than the tolerance.

### Simulation

#### Individual Foci

First, a single actin polymerization focus is simulated using the Monte Carlo model (Sec. Actin dynamics at individual foci) with fixed ‖**F**_*mem*_‖ in Eq. (2). The focus is initialized with different numbers of barbed ends (between 1 and 20) and simulated until all barbed ends had vanished. Hereby, the number of barbed ends in each time step as well the lifetime of the focus were tracked. In order to compare the outcomes of these simulations with theoretical expectations, we also derive the rate equations for this simple system. Assuming that a focus has *B* filaments with barbed ends, which divide into *m*_*c*_ filaments with capped and *m*_*u*_ filaments with uncapped minus ends, we obtain

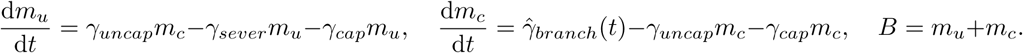

As 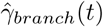 in turn depends on *B*, these equations are highly nonlinear and have been solved for their stationary state numerically.

#### 2D Model

Simulations are performed in MATLAB on a desktop computer. Table 1 contains the parameters used in the simulations, unless stated otherwise. We first initialize the mesh by tracing a circle with equidistant vertices of *δ*_*s*_, then the vertices of PSD (neck) as described in Section Membrane mesh initialization and morphological constraints. We then simulate an initialization period in which the mesh points **x**^*k*^ move considering only the force generated by the membrane. Thus, during this initialization period the spine shape shrinks until it reaches a stable configuration, which we refer to as the resting shape.

As discussed above, the finite elements approximations of the geometrical properties are only valid when the mesh is dense enough. If the vertices move too far apart from each other these properties are lost, and therefore, the mesh has to be redefined. Thus, we perform remeshing at each timestep by calculating the distance between two neighbouring vertices and remove one if the distance between them is below *d*_*min*_ = (3*/*5)*δ*_*s*_ or add a new vertex in between if the distance is above *d*_*max*_ = (4*/*3)*δ*_*s*_.

After finding the resting shape configuration of the dendritic spine, we include actin dynamics and forces in the simulation (Sec. Actin dynamics at individual foci and Sec. Actin-generated force). For this, initially, 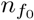 actin polymerization foci are inserted as described in Section Foci generation and removal and the generation of new foci is enabled. Note, the indicated simulation times start after the initialization phase. During the simulations (Fig. 3) we track the spine shape by saving the mesh regularly as well as the spine area, which is recorded every time step. To assess the influence of different model parameters (Fig. 4-10), we perform one 90 minute simulation for each parameter value and determine the mean, standard deviation and auto-correlation function of the spine area fluctuations. Moreover, we evaluate the distribution over the assumed values of the number of foci and barbed ends and the mean lifetime of polymerization foci. We then perform fifteen 15-minute-simulations to obtain statistics for different initial conditions, estimate their uncertainty, and test whether values vary significantly.

**Fig 5.**
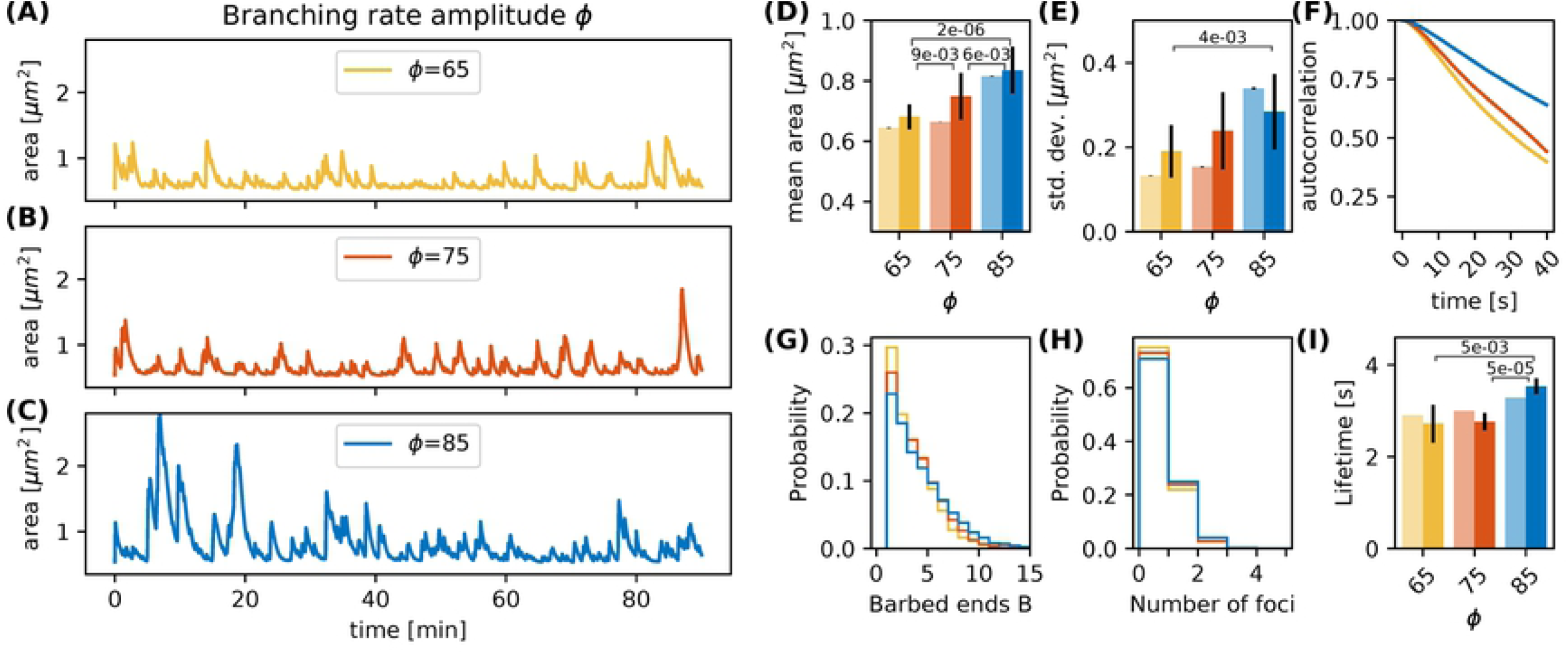
Varying the branching rate amplitude *ϕ*. (A-C) Evolution of the spine head area over time for different values of *ϕ*: (A) *ϕ* = 65 (B) *ϕ* = 75 and (C) *ϕ* = 65. (D) Temporal mean of the spine head area. Light bars correspond to values from single 90 minute simulations (standard deviations obtained from 50-fold bootstrap). Full colored bars and errors correspond to the mean and standard deviation over fifteen 15 minute simulations. The p-value for significant Welch-tests is indicated. (E) Standard deviation of the spine head area over time. (F) Autocorrelation functions for the area fluctuations in the 90 minute simulations. (G) Relative frequency of the mean number of barbed ends per focus over all simulation time-steps from panels A-C. (H) Relative frequency of the number of actin polymerization foci. (I) Mean lifetime of a focus.

**Fig 6.**
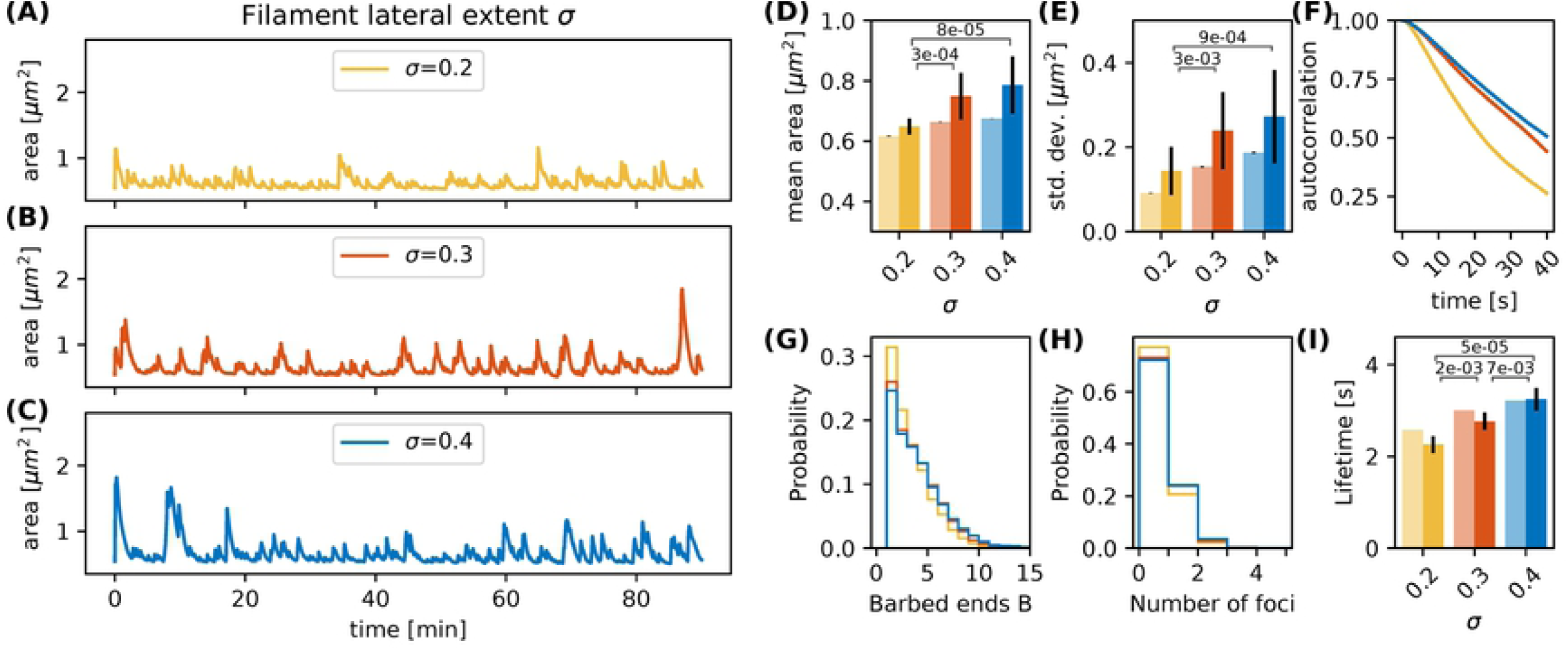
Varying the filament lateral extent *σ*. (A-C) Evolution of the spine head area over time for different values of *σ*: (A) *σ* = 0.2 (B) *σ* = 0.3 and (C) *σ* = 0.4. (D) Temporal mean of the spine head area. Light bars correspond to values from single 90 minute simulations (standard deviations obtained from 50-fold bootstrap). Full colored bars and errors correspond to the mean and standard deviation over fifteen 15 minute simulations. The p-value for significant Welch-tests is indicated. (E) Standard deviation of the spine head area over time. (F) Autocorrelation functions for the area fluctuations in the 90 minute simulations. (G) Relative frequency of the mean number of barbed ends per focus over all simulation time-steps in panels A-C. (H) Relative frequency of the number of actin polymerization foci. (I) Mean lifetime of a focus.

**Fig 7.**
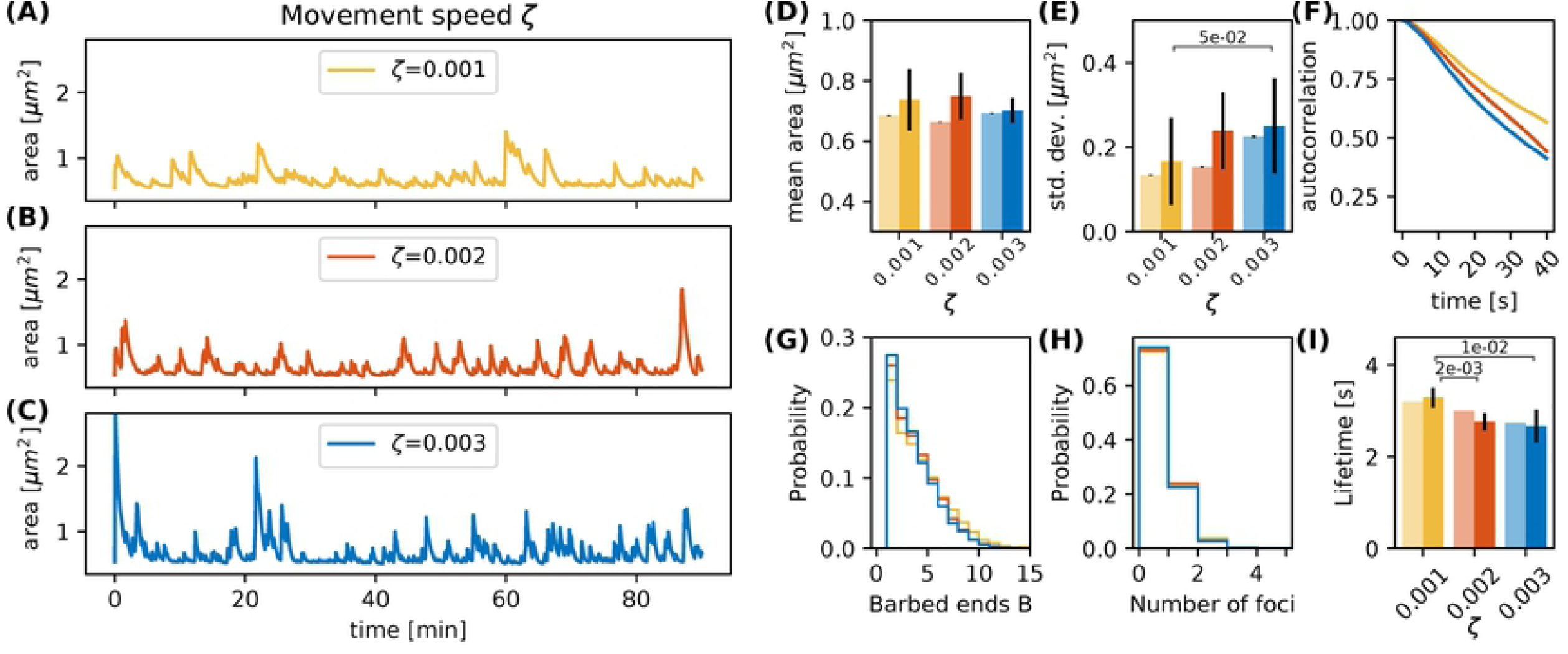
Varying the movement speed *ζ*. (A-C) Evolution of the spine head area over time for different values of *ζ*: (A) *ζ* = 0.001 (B) *ζ* = 0.002 and (C) *ζ* = 0.003. (D) Temporal mean of the spine head area. Light bars correspond to values from single 90 minute simulations (standard deviations obtained from 50-fold bootstrap). Full colored bars and errors correspond to the mean and standard deviation over fifteen 15 minute simulations. (E) Standard deviation of the spine head area over time. The p-value for significant Welch-tests is indicated. (F) Autocorrelation functions for the area fluctuations in the 90 minute simulations. (G) Relative frequency of the mean number of barbed ends per focus over all simulation time-steps in panels A-C. (H) Relative frequency of the number of actin polymerization foci. (I) Mean lifetime of a focus.

**Fig 8.**
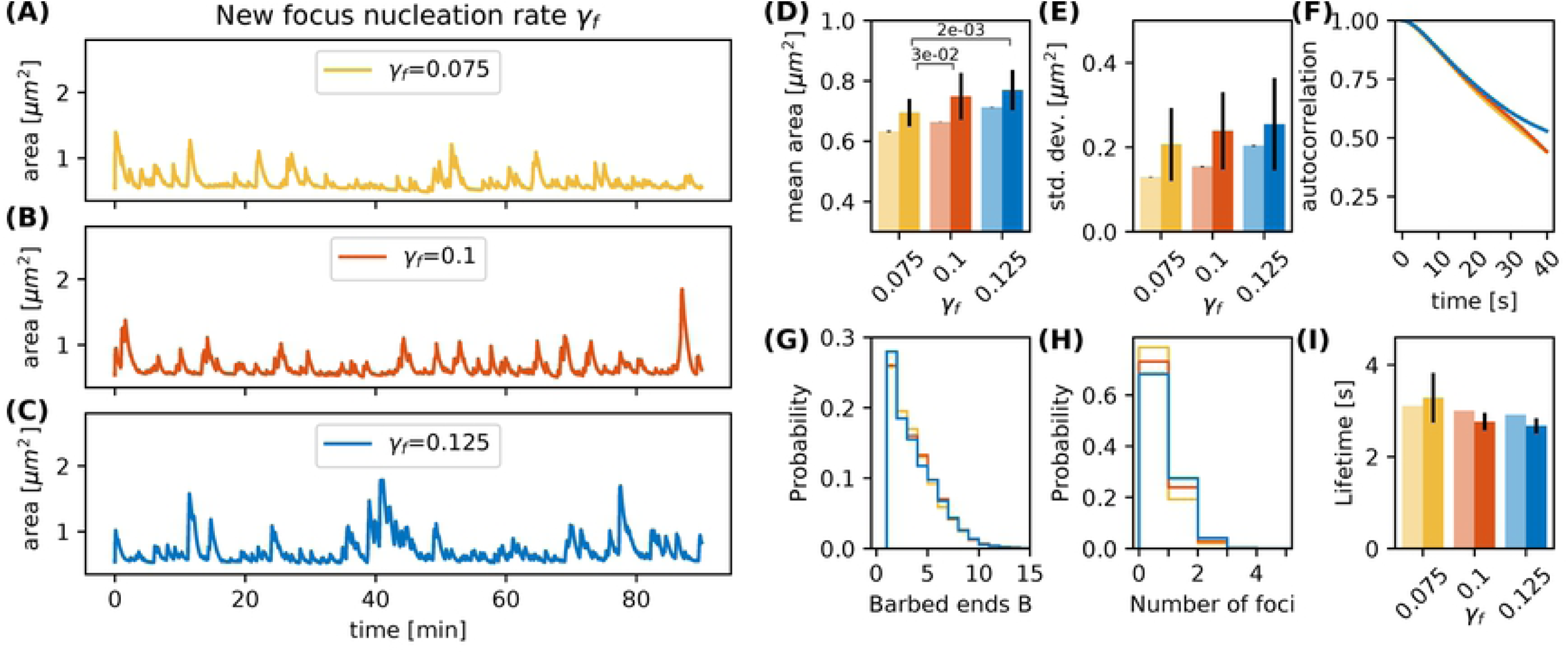
Varying the focus nucleation rate *γ*_*f*_. (A-C) Evolution of the spine head area over time for different values of *γ*_*f*_ : (A) *γ*_*f*_ = 0.075 (B) *γ*_*f*_ = 0.100 and (C) *γ*_*f*_ = 0.125. (D) Temporal mean of the spine head area. Light bars correspond to values from single 90 minute simulations (standard deviations obtained from 50-fold bootstrap). Full colored bars and errors correspond to the mean and standard deviation over fifteen 15 minute simulations. The p-value for significant Welch-tests is indicated. (E) Standard deviation of the spine head area over time. (F) Autocorrelation functions for the area fluctuations in the 90 minute simulations. (G) Relative frequency of the mean number of barbed ends per focus over all simulation time-steps in panels A-C. (H) Relative frequency of the number of actin polymerization foci. (I) Mean lifetime of a focus.

**Fig 9.**
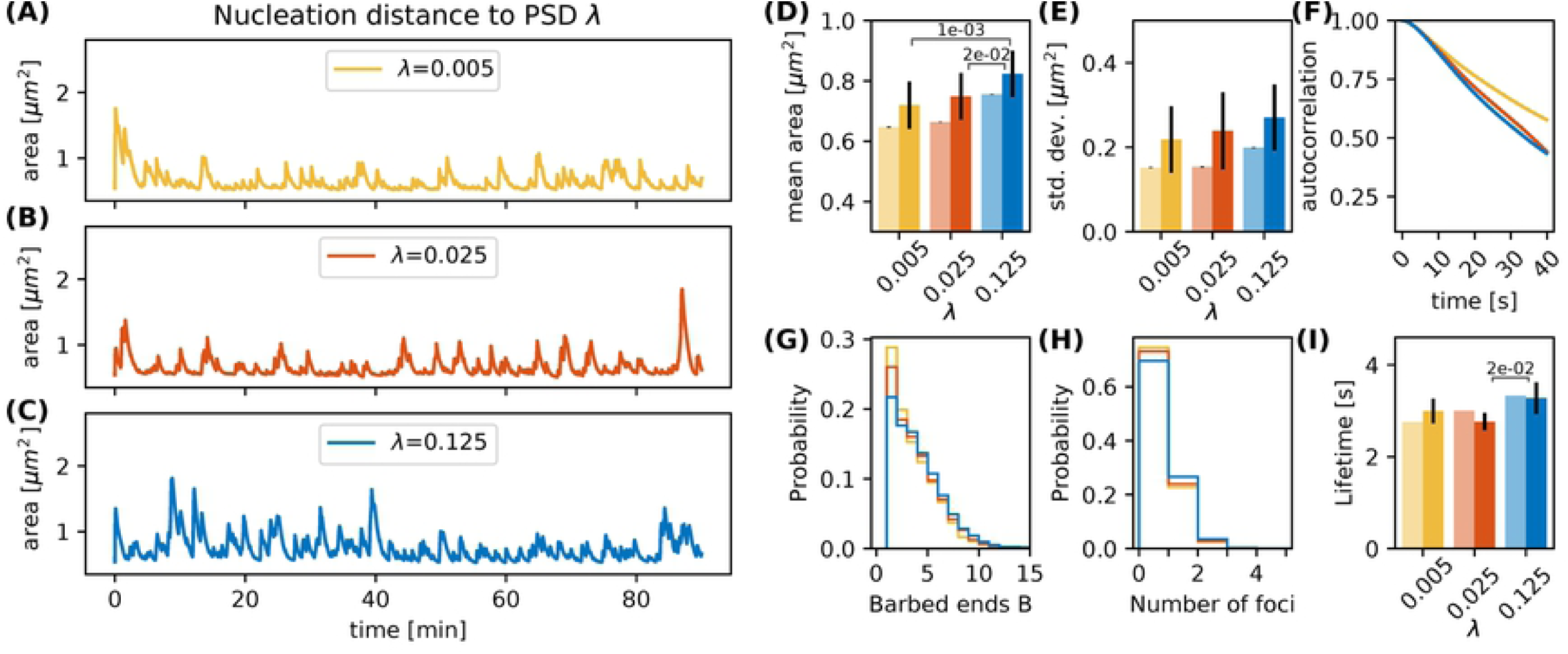
Varying the nucleation distance parameter *λ*. (A-C) Evolution of the spine head area over time for different values of *λ*: (A) *λ* = 0.005 (B) *λ* = 0.025 and (C) *λ* = 0.125. (D) Temporal mean of the spine head area. Light bars correspond to values from single 90 minute simulations (standard deviations obtained from 50-fold bootstrap). Full colored bars and errors correspond to the mean and standard deviation over fifteen 15 minute simulations. (E) Standard deviation of the spine head area over time. The p-value for significant Welch-tests is indicated. (F) Autocorrelation functions for the area fluctuations in the 90 minute simulations. (G) Relative frequency of the mean number of barbed ends per focus over all simulation time-steps in panels A-C. (H) Relative frequency of the number of actin polymerization foci. (I) Mean lifetime of a focus.

**Fig 10.**
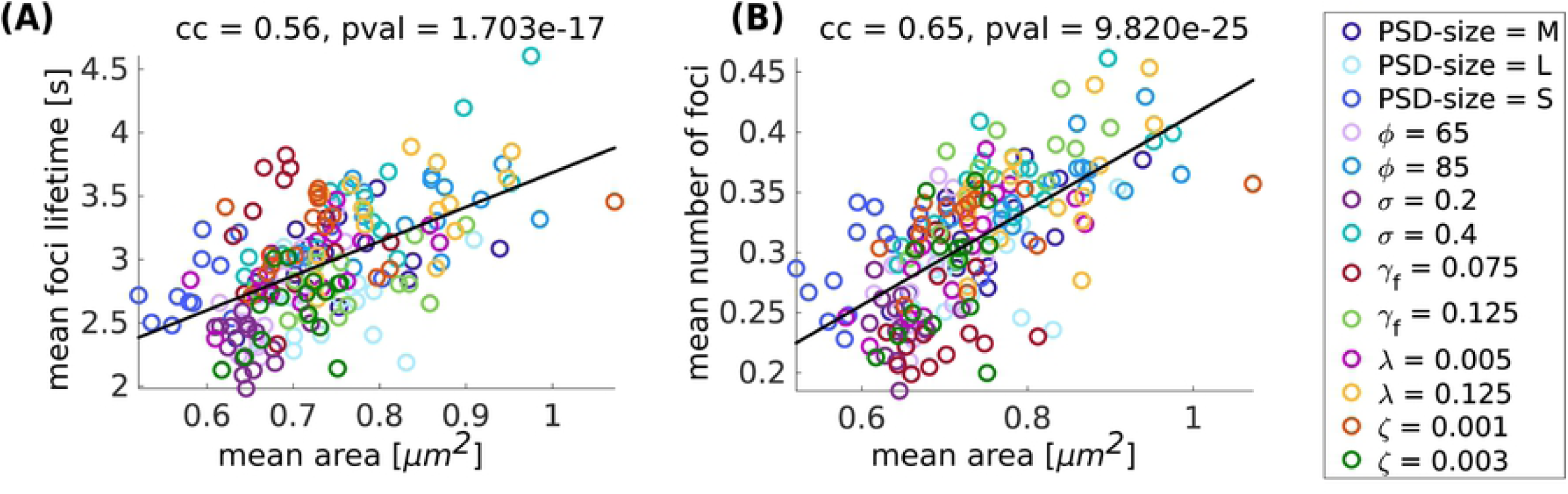
Summary of the parameter variation simulations. (A) Mean focus lifetime over mean spine area for different parameter variations. Each dot represents one simulation with a parameter variation corresponding to its color. Standard values of the parameters are reported in Table 1. For every variation of a parameter 15 simulations of 15 minutes were performed. cc denotes the linear correlation coefficient and pval is the p-value for testing the hypothesis of no correlation against the alternative hypothesis of a nonzero correlation, using Pearson’s Linear correlation coefficient. (B) Same for mean number of foci over mean area.

#### 3D Model

Simulations in 3D are performed as in 2D. But in this case, the mesh is initialized using the MATLAB function icosphere.m [36] which generates a unit geodesic sphere that we multiply by *r*_*s*_. Additionally, in 3D the mesh has to be isotropic and conserve the number of neighbours of each vertex at each timestep to maintain the geometrical properties of the finite elements approximation. Thus, remeshing is implemented using the MATLAB function remeshing.m [37], that is based on OpenMesh [38, 39]. The target edge length is set to *δ*_*s*_ and three iterations are performed each time. To compare the 3D-shapes with the 2D simulations we perform a two dimensional projection of the three dimensional mesh. For this, we project all points to the x-z-plane and trace a boundary around the projected points using the MATLAB function boundary.m. We do the same for another projection to the y-z-plane to compare the two 2D projections of the same spine.

## Results

### Individual polymerization foci have finite lifetime depending on the membrane force

In order to understand the highly nonlinear interaction between actin dynamics and membrane force, we first simulated the model for a single actin polymerization focus with a constant counteracting membrane force **F**_*mem*_ in Eq. (2). We then tracked the time course of the number of barbed ends in our Monte-Carlo model (Fig. 2A) and find that it fluctuates around a mean value (dashed lines) on a timescale of seconds. This mean value depends on the counteracting force. If **F**_*mem*_ increases, the mean number of barbed ends decreases (blue and orange dashed lines in Fig. 2A, Fig. 2B-C).

Importantly, the foci have nonzero probability to transit to *B* = 0. At that point no barbed end can be generated by branching anymore and the focus is dynamically dead. As a result all foci have limited lifetime. We also tracked this lifetime at different counteracting forces and find that it decreases when increasing the force (see Fig. 2D). We finally evaluated the relation between the mean number of foci and the lifetime for varying forces and find, as expected from the above reasoning, that foci with more barbed ends live longer (Fig. 2E).

The dependency of mean number of barbed ends (Fig. 2C) and lifetime of a focus (Fig. 2D) on the membrane force can be explained by the fact that increasing **F**_*mem*_ decreases the treadmilling velocity *v*_*T*_; and hence, the branching rate (see Eq. 2). Accordingly, the mean number of barbed ends is smaller and the distribution is shifted towards smaller numbers of *B* (Fig. 2B-C). This increases the probability of being at *B* = 1, and, in turn, the probability to reach zero barbed ends. Consequently, the lifetime of these actin polymerization foci decreases when increasing the counteracting force. This relation between membrane force and focus lifetime indicates that the spine shape, which determines the membrane force, influences magnitude and the temporal properties of the shape fluctuations.

### Shape-fluctuations of the spine

In the next step, we studied the interplay between the membrane and actin forces. For this, we simulated the full model with multiple actin polymerization foci distributed within a spine head (Figs. 3A-B). We initialized the model with four of such foci, that push the spine membrane outward. However, as the lifetimes of foci (Fig. 3C and G) are much shorter than the lifetime of spines, which can persist over months [40], there must be a focus creation mechanism (i.e., nucleation of new actin polymerization foci) Without such a nucleation mechanism all foci quickly reach zero barbed ends (in less than 9 seconds in Fig. 3G) and the spine returns to the resting shape (gray line in Fig. 3B). Moreover, also spine area fluctuations cease (gray line in Fig. 3D). We therefore included a foci creation mechanism where new foci are created at the beginning of each time step at a rate *γ*_*f*_. As it is not clear where such nucleations happen, we also introduced a distance parameter *λ* which allows us to scale continuously between focus nucleation everywhere within the spine and nucleation close to the PSD, as suggested by [19]. The influence of this parameter on the emerging shape fluctuations will be investigated in detail later (see Sec. Nucleation rate *γ*_*f*_ and location *λ*).

Figures 3 A and B (blue line) show the resulting shape dynamics of the spine in our model. The proposed nucleation mechanism together with the short lifetimes of individual foci allows the spine to have different asymmetric shapes over time, which are qualitative similar to the experiments [27]. Note that during the depicted time interval, several foci have died out and several others have been nucleated (Fig. 3E), which also induces major fluctuations in the number of barbed ends (Fig. 3F). In general, we observed that spine area increases when several foci are active at the the same time or when a focus is long-lived (Figs. 3D-E). Thus, shape and area fluctuations of a spine are the result of the transiently active foci working against the membrane. In particular, they are generated by the stochasticity of the molecular dynamics of actin filament assembly, which eventually leads to the die-out of a focus. Therefore, it can be expected that these dynamics as well as the mechanics through which they interact with the membrane (see previous section) will have a major impact on the emerging fluctuation.

### Influence of model parameters

In order to better understand how spine size fluctuations are affected by the dynamics of actin and the interplay between forces generated by actin polymerization foci and spine membrane, we investigated the effect of varying multiple model parameters. For this, we used the parameters in Table 1 and increased or decreased the value of one selected parameter at a time.

#### Size of the postsynaptic densitiy

First, the spine size was varied by changing the size of the PSD (*r*_*PSD*_, see Sec. Membrane mesh initialization and morphological constraints). If the radius of the PSD is enlarged to *r*_*PSD*_ = 0.4330, this also affects the distance between PSD and neck and alters also the resting shape of the spine (Fig. 4A-C, black line). Accordingly, the mean spine area, evaluated over 90 minute simulations of individual spines, increases with the PSD size (Fig. 4G, pale bars). To test whether this tendency is significant, we performed a Welch-test comparing the mean spine areas in fifteen 15 minute simulations for each PSD size (Fig. 4G, full colored bars). We find that at least the small PSD size (set to *r*_*PSD*_ = 0.2179) gives rise to significantly smaller mean areas. Moreover, although the medium PSD-size spine from the 90 minute simulation shows a smaller area standard deviation than the 90 minute simulations of spines with different PSD-size (Fig. 4H, pale bars), multiple simulations show that the area standard deviations are similar. We also evaluated how quickly the autocorrelation function of the area fluctuation decays (Fig. 4I). Here, we found that the area of spines with large PSD-size is temporally longer correlated to itself, indicating that the fluctuations occur more slowly. One reason for this might be that the membrane forces typically decrease for larger spines such that the decay back to the rest shape happens more slowly. Interestingly, the autocorrelation for spines with medium PSD size decays even faster than that for small PSD size. This may be due to the fact that the actin polymerization foci in spines with smaller PSD tend to last longer (Fig. 4L).

The 90 minute simulation of the spine with a large PSD-size also exhibits a period (between minutes 60 and 70) where the spine area fluctuations cease (blue bar in Fig. 4F, spine shapes in Fig. 4C.II). Although new foci are nucleated during this period, changes in spine shape are minor because new foci randomly nucleate nearby each other (locations marked by circles in Fig. 4C.II). Moreover, the curvature near these foci is large, such that the force generated by the membrane is higher decreasing the branching rate; and hence, the lifetime of those foci.

#### Branching rate amplitude *ϕ*

The branching rate of actin polymerization foci *γ*_*branch*_ in Eq. (2) is scaled by an amplitude *ϕ*. Three simulations with different values of *ϕ* over 90 minutes, as well as the fifteen 15 minutes repetitions with different initial conditions, show that an increase of *ϕ* enlarges the mean spine head area significantly (Fig. 5D, p-value for significant Welch-tests indicated). Although not in the same proportion, a decrease of *ϕ* reduces the spine area (Fig. 5A and D). This relation between the mean area and *ϕ* can be explained by the fact that spines with a larger value of *ϕ* also have more barbed ends at the actin polymerization foci (Fig. 5G). Due to this increased number of barbed ends, the polymerization foci of those spines tend to last longer (Fig. 5I). As a consequence, there are more actin polymerization foci for spines with a larger branching rate amplitude (Fig. 5 H), which push the membrane outwards and increase the area. A similar picture emerges for the magnitude of the fluctuations measured by the standard deviation (Fig. 5E) of the area as well as for the timescale of the autocorrelation decay (Fig. 5F). Especially for large values of *ϕ* we observe a significantly larger standard deviation (Fig. 5E) and a slower autocorrelation decay (Fig. 5F). This fits well with the idea that the polymerization foci are more long lived; and therefore, push the membrane outward for longer times leading to larger area deviations. We conclude that an increase of *ϕ* enlarges the mean, the standard deviation and the autocorrelation decay timescale of the spine area fluctuations due to an increase of the lifetime of actin polymerization foci. However, a decrease in *ϕ* by the same magnitude does not affect the spine area to the same degree, which highlights that the underlying processes are subject to nonlinear interactions.

#### Lateral extend of actin filament *σ*

Besides the number of barbed ends at the actin polymerization foci, the actin generated forces on the membrane and the resulting deformations of the membrane are also determined by the lateral spatial extent of actin filaments (*σ* in Eq. (3)). When the lateral extent is small, the width of the bump that a focus causes in the membrane is narrow. The shape of this bump has a direct effect on the geometrical properties of the spine membrane around the focus. For example, a narrow protrusion has a greater curvature, which produces an increase in ‖**F**_*mem*_‖ working against this deformation. This entails a decrease in the branching rate as well as in the number of barbed ends (Fig. 6G) and leads to less active foci with a shorter lifetime (Fig. 6H and I). This shorter lifetime of foci also implies that foci push the membrane for shorter time such that the variations in the spine area become smaller (Fig. 6E) and decay faster (Fig. 6F).

#### Movement speed *ζ*

The conversion factor between force imbalance and movement *ζ* can be expected to have a strong influence on the magnitude of spine shape change per time step. Judging from the dynamics shown in Figure 7A-C, the area fluctuations also seem to be much faster when increasing *ζ*. However, this is mostly due to an increase in the amplitude of the fluctuations (Fig. 7E) whereas the timescale of the autocorrelation decay remains relatively constant. Note as an increase of *ζ* enhances spine fluctuation extent, it also affects the membrane geometrical properties, the membrane force and, hence, the barbed end branching rate. This leads to less barbed ends (Fig. 7G) and a reduction of the foci lifetime (Fig. 7I). Still, in sum, spines with different values of *ζ* have similar mean area over time (Fig. 7 D).

#### Nucleation rate *γ*_*f*_ and location *λ*

Besides the parameters that influence force generation and translation to movement, also the parameters of the nucleation mechanism proposed in Section Shape-fluctuations of the spine can have a strong influence. First, we vary the nucleation rate *γ*_*f*_ at an intermediate value of the PSD distance scaling parameter *λ*. As expected, an increase in *γ*_*f*_ raises the number of actin polymerization foci and the spine area over time (Fig. 8H). This leads to a significant increase in the mean area and a trend towards increasing standard deviations (Fig. 8D-E). Although these foci have slightly shorter lifetimes (Fig. 8I), the decay of the autocorrelation remains at the same timescale (Fig. 8F). The main reason for the reduction of foci lifetime is the feedback between the number of barbed ends and the branching rate in Eq. (2). If *B* increases then *γ*_*branch*_ decreases ensuring a limited number of barbed ends at the actin polymerization foci.

The location for the polymerization of new foci depends on the distance from the PSD scaled by parameter *λ*, as stated in Section Simulation. For larger values of *λ*, the nucleation points are more likely located more distant to the PSD and the spine mean area is larger due an increase in the lifetime of the actin foci (Figs. 9 E-I). We speculate that this can be explained by the fact that for small *λ* all foci nucleate close to the PSD. Hence, all foci push outward the same small fraction of the membrane, which thereby assumes a strong curvature. This, in turn, leads to a strong counteracting force and hence a shorter lifetime of the foci.

In conclusion, we find that geometrical constrains as well as parameters related to actin filament assembly, force generation and focus nucleation have a strong influence on the emerging fluctuation. We summarized the most prominent effects in Table 2.

**Table 2.**
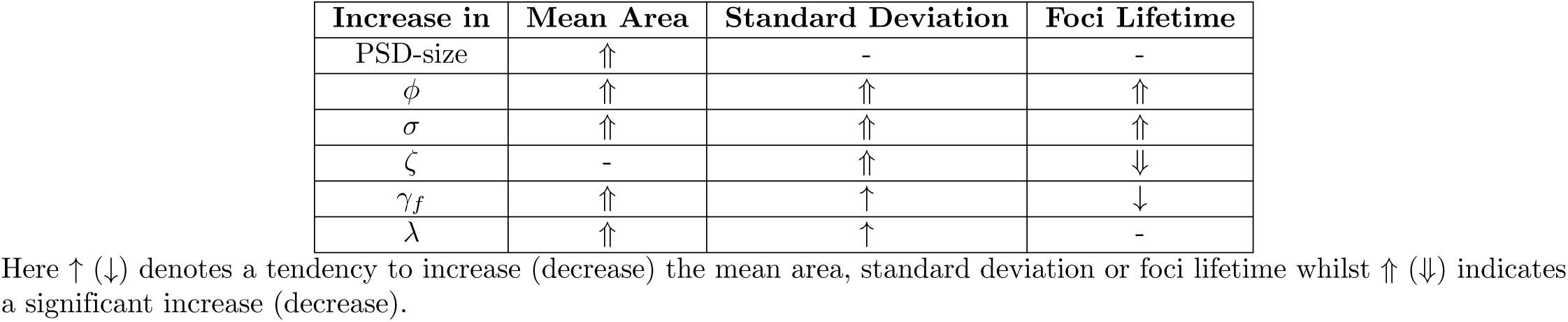
Summary Table.

#### Influence of parameter variation on spine area

After evaluating the influence of the individual parameters, we investigated whether there are general relations between the evaluated quantities that are preserved over all these variations. To investigate this, we used the fifteen 15 minute simulations for each parameter variation and plotted the values of mean area, focus lifetime and mean number of foci for each of these individual simulations against each other. On the one hand, we find that spines with greater mean area over time, have larger mean foci lifetimes (Fig. 10A). However, spines with smaller mean area can also have long-lasting foci when the force generated by the membrane is not affecting the branching rate strongly. For example, when the focus nucleation rate *γ*_*f*_ is high or the movement speed *ζ* is small. On the other hand, there is a positive correlation between the mean number of actin polymerization foci and spine mean area (10B), which has also been found in experimental data from [19]. These results imply that the macroscopic spine area fluctuation is heavily relying on the stochastic dynamics of the actin polymerization foci and filament dynamics therein.

### Correlation of the number of foci and spine area fluctuations

Although the spine shape is determined by a complex interplay between forces emerging from actin activity and the geometrical properties of the membrane, the above described correlations indicate that there is a strong link between spine area and its polymerization foci. Therefore we investigated whether the number of polymerization foci at each time-step can be used to predict not only the mean but also the time-course of the spine head volume/area, which is commonly measured in experiments. As the expanding force in our model comes from the actin foci, we first tested whether there is a relationship between number of actin polymerization foci and the spine head area. To quantify this, we tracked the area and the number of foci throughout a 90 minute simulation of a spine (Fig. 11A) and evaluated the correlation between these quantities. We found a significant correlation, but with a very small correlation coefficient (Fig. 11B). When examining the time courses in Figure 11A, we see that when there is no focus the area shrinks to a state close to the resting shape area and a slight increase in area when the number of foci increases. Hence, we also investigated the relationship between the actin foci and spine area changes Δarea (Fig. 11C) and found that there is indeed a significant correlation with a high correlation coefficient between these quantities. Thus, we constructed a simple model that predicts the area of a spine using only the number of foci at a given time-step. Apart from the area change being proportional to the number of foci, we also included a mean retrieval that drives the area back to the area of the Helfrich resting shape area *A*_*s*_. In particular, an estimator *Ā* for the spine area *A* at each time step *t*_*j*_ can be recursively calculated by the following model

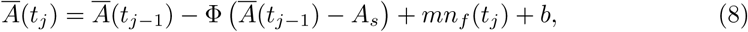

where the term *mn*_*f*_ (*t*_*j*_) + *b* accounts for the change of area that scales linearly with the number of actin polymerization foci *n*_*f*_ at time *t*_*j*_ and Φ represents a decay rate to the resting area *A*_*s*_, which we extracted from our simulations. The model parameters *m, b* and Φ from Eq. (8) and the initial area *Ā*(*t*_0_) were fitted using the nonlinear least square method and the area trace of Figure 11A from minute 1 to 60 (Fig. 11 D, fit results: root mean square error (RMSE) = 0.0562,

**Fig 11.**
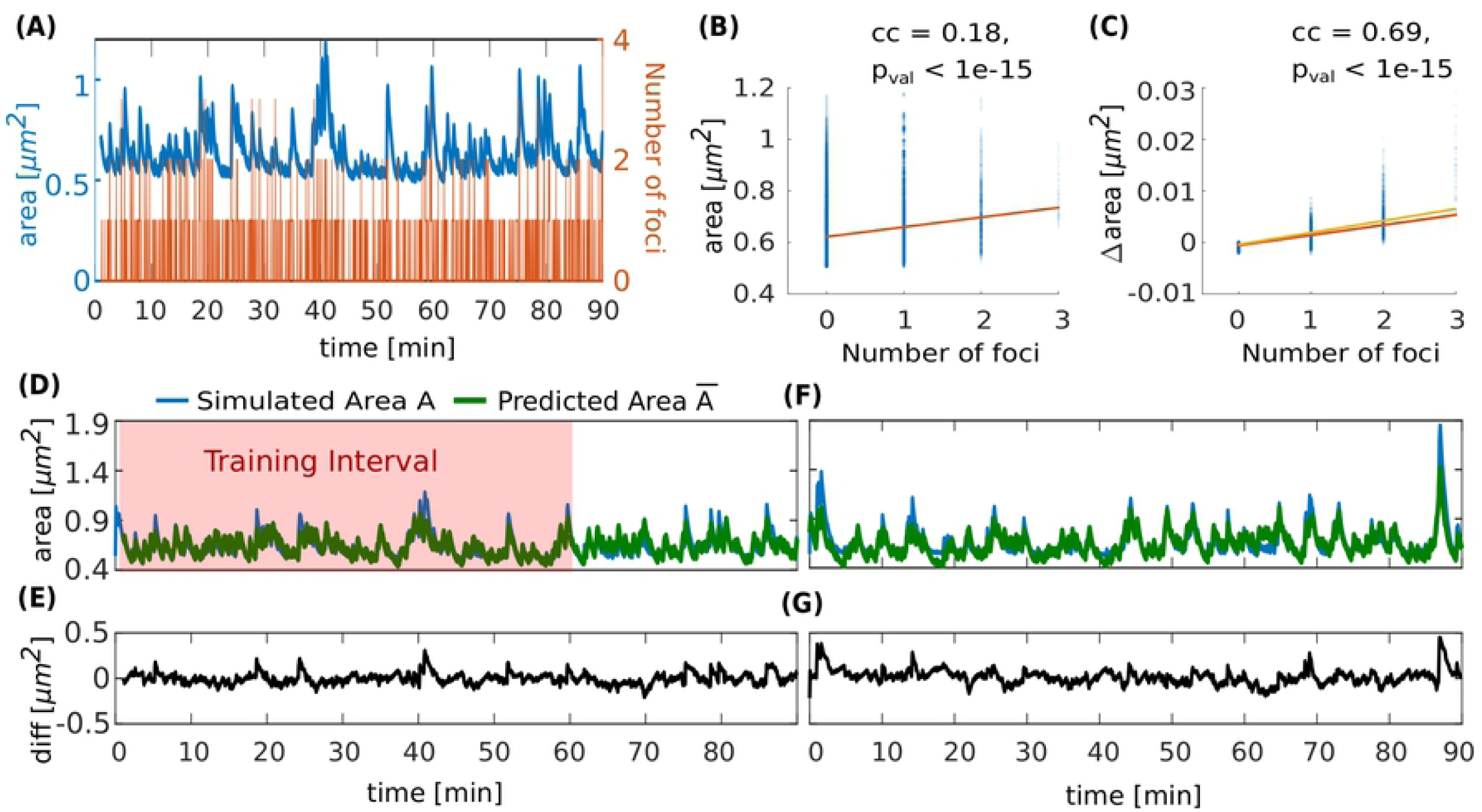
Temporal evolution of spine area can be predicted by number of actin polymerization foci. (A) Evolution of number of actin polymerization foci (orange line) and spine area (blue line). (B) Spine area over the number of actin polymerization foci for each time-step of the simulation. Color saturation indicates overlay of data points; darker regions contain more data observations. Orange line is a linear fit to the data, cc denotes the linear correlation coefficient and pval is the p-value for testing the hypothesis of no correlation against the alternative hypothesis of a nonzero correlation, using Pearson’s Linear Correlation Coefficient.(C) Same for the difference in spine area over the number of actin polymerization foci. Yellow line is a linear function using the parameters fitted to Eq. (8) in panel D. (D) Simulated area (blue line) and its approximation by estimator *Ā* (green line, Eq. (8)) over time. Red shaded zone depicts the training data set for fitting the estimator parameters. (E) Difference between simulated and predicted area in (D). (F) Simulated area (blue line) corresponding to the plot in Figure 5B and its approximation (green line) over time. (G) Difference between the simulated and predicted area in (F).

*Ā*(*t*_0_) = 0.7507, *m* = 0.002734, *b* = *-*0.0004135, Φ = 0.002734). Hereby, the obtained values for *b* and *m* are close to a linear fit to the relation between the number of foci and the change of the area (orange line in Fig. 11C; Δarea = *m*′*n*_*f*_ + *b*′, with *m*′ = 0.00197 and *b*′ = *-*0.000570). Also, and *Ā*(*t*_0_) is close to the actual start value *A*(*t*_0_) = 0.7239. Given that our area estimator is recursive and could accumulate errors over time, *Ā* performs well, even for a time interval it was not fitted to (Fig. 11D from minute 60 to 90, RMSE = 0.0652). Moreover, it could even be applied to a different simulation with the same parameters (Fig. 11F, RMSE = 0.0822). Note that the estimator error increases in periods with large areas (Fig. 11E,G), which may be due to the fact that the relation between foci and the change in area may be nonlinear (compare Fig. 11C). Nevertheless, we can conclude that area fluctuations can be predicted very well from the number of actin polymerization foci. This again underlines a strong link between the microscopic stochastic dynamics at the actin polymerization foci and the macroscopic area fluctuations.

### Model extension to 3D

So far we have only considered spine shapes in 2D, but in order to verify if the dynamics of actin polymerization foci influence spine shape fluctuations in real spines in a similar way, we extended our model to 3D. In this extended model, actin dynamics are preserved but the membrane mesh, all positions and forces are adapted to 3D (see S3 Appendix).

To check whether the 3D and the 2D simulation are comparable, we assumed that the 3D spine shape is observed as in a microscope and projected to a two dimensional plane. Hereby, we perform projections from multiple sides, in particular to the y-z- and the x-z-plane (Fig. 12B-C and G-H, respectively). We find that the fluctuations from the projected 3D model are similar to the area fluctuations from the 2D model when adjusting the branching rate amplitude *ϕ* and movement speed *ζ* (Fig. 12D). Note that deformations in the 3D model where the acting forces are not in the projection plane will appear to be effectively slowed down in 2D projections. To compensate for this, the movement speed of the 2D model has been slowed down by adjusting *ζ*. Moreover, multiple foci in the the 3D model may be projected onto the same 2D bump such that it appears as if the foci were more long-lived than they are. The adjustment in *ϕ* allows the 2D model to have more barbed ends such that the foci actually last longer. After adjusting the parameters for these geometrical properties, 3D to 2D simulations exhibit qualitatively similar fluctuations.

**Fig 12.**
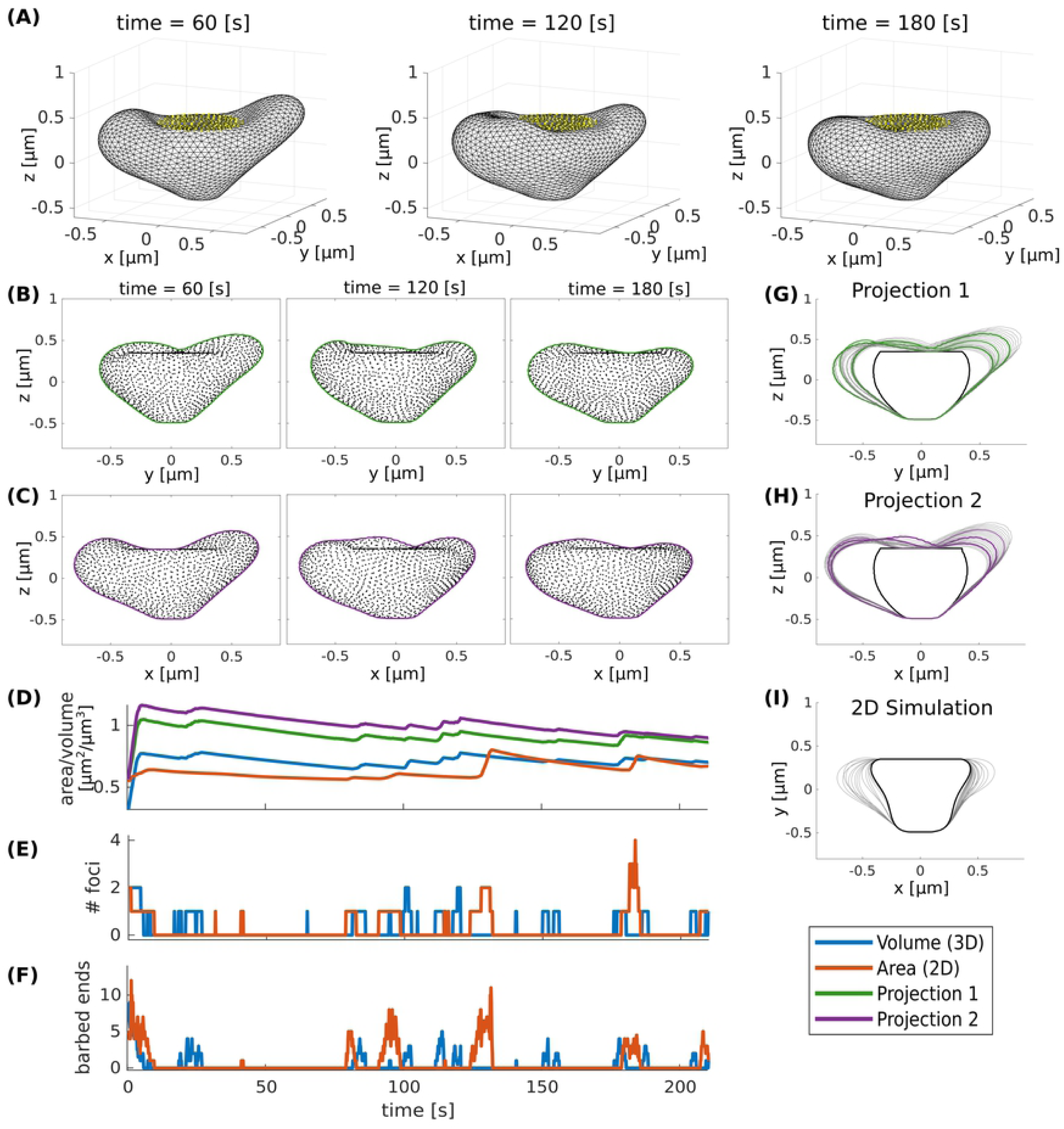
Comparison between 3D and 2D model. (A) 3D simulation snapshots at different time points. Yellow dots indicat the PSD. (B-C) Projection of the 3D simulation to two 2D views. Black dots are the corresponding 3D vertices and lines are a fitted boundary around these points. (D) Time evolution of volume (3D simulation) and area (2D simulation and 2D projections). (E) Number of actin foci during the 2D/3D simulation. (F) Mean number of barbed ends. (G-H) Spine shapes emerging from the 2D projections of the 3D model in time intervals of 10 seconds. Black lines represent the initial shape and green and purple lines correspond to the snapshots in (B) and (C), respectively. (I) Spine shapes emerging from 2D simulation of the model. Parameters for 3D and 2D simulations are in Table 1 except that in 3D *δ*_*s*_ = 0.06, *ζ* = 0.004 and *ϕ* = 30, and in 2D *ζ* = 0.001.

## Discussion

We proposed a model for dendritic spine shape fluctuations based on actin polymerization foci. The model was used to simulate the dynamics of a single focus as well as the interaction of multiple foci within a spine in 2D and 3D and the resulting fluctuations of the spine head shape were analyzed. Variations of the model parameters revealed that the properties of the molecular processes and mechanics have strong influence on the emergent shape fluctuations. Along these lines, strikingly, our model predicts that the fluctuations of the spine head size could be very well predicted from only knowing how many foci were active over time. Thus, our model provides a platform to study the relation between molecular and morphological properties of the spine.

The proposed model is, to our knowledge, one of the first to reproduce the rapid asymmetric shape fluctuations (Fig. 3) observed in experiments [27]. These asymmetric shape fluctuations result from local imbalances between forces generated by membrane deformation and forces generated by the barbed ends in active actin polymerization foci. Strikingly, these foci have a limited lifetime due to the stochastic nature of the actin filament dynamics. Thereby, the stochasticity of actin dynamics is also transferred onto the spine shape and size, which is evidenced by the fact that the number of active foci can predict the spine area (Figs. 10-11). Our model predicts that the focus lifetime is inversely proportional to the force generated by the spine membrane (Fig. 2), which caused by a feedback between this force and the branching rate. This mechanism, thus, couples geometric properties and molecular dynamics, and links the dynamics of multiple foci via the membrane.

Due the limited lifetime of the actin polymerization foci, we proposed a nucleation mechanism due to which foci are stochastically generated at different locations in the spine. The generation rate and the initial location of these new foci have a great impact on the evolution of spine area over time. For example, an increase of the nucleation rate causes increases of the mean spine area and its standard deviation (Fig. 8).

Interestingly, foci generated with fast nucleation rate also tend to have a shorter lifetime evidencing a saturating mechanism or self-regulation (compare [13]). Moreover, we used our model to test the influence of the nucleation location of these foci. Experimentally, it has been observed that actin foci are mainly located at the tip of the spine [18, 19] and the branching protein Arp2/3, which is necessary to build branched actin filaments, is mainly located in a doughnut-shaped zone around the PSD [41]. Such a constraint on the nucleation location of polymerization foci also has a strong impact on the shape fluctuations of our model-spines (Fig. 9): When foci nucleate closer to the PSD, they tend to last for shorter time intervals such that the mean number of foci is smaller which, in turn, reduces the mean area of the spine. This demonstrates that changes in the polymerization activity can be caused only by differences in geometry without changing any reaction rates..

Furthermore, we observed that, despite the change of shape, the spine area always fluctuates around a mean value, in agreement with experimentally observed spine fluctuations on short timescales [27]. However, this mean value, as well as the magnitude and timescale of the fluctuations are affected by various model parameters. For example, there is a strong influence of the PSD-size on the mean spine area (Fig. 4) which is in line with the experimentally observed correlation between these quantities [42, 43]. Similarly, reducing the branching rate in our model by decreasing *ϕ* leads to a decrease in the mean and standard deviation of spine area (Fig. 5), which is in line with findings that the branching factor Arp2/3 is necessary for spine enlargement and maintenance of spine morphology [44]. Furthermore, an increase of the movement speed parameter *ζ* leads to a increase in spine area standard deviation (Fig. 7), which has been similarly observed experiments that artificially decreased the density of the extra-cellular matrix in visual cortex [45]. Overall, these results indicate that the mean spine size as well as the magnitude and timescale of spine shape fluctuations are regulated by the properties of the underlying molecular processes (e.g., reaction rates, force generation). Therefore, our model can represent a broad variety of different fluctuation characteristics as observed in experiments through different parametrizations.

In the future, the model can be extended in various directions: On the one hand, the shape fluctuations may, on longer timescales, influence the model parameters such as PSD size, molecule concentrations and reaction rates. Hence, the mean area around which the spine fluctuates as well as other fluctuation characteristics could be continuously adapted giving rise to a slower feedback-loop (compare [9] and [13]). On the other hand, so far, the proposed model only considers spines at basic neuronal activity. However, in the future, it can be extended to include induction of activity-dependent plasticity (LTP/LTD) by explicitely modelling the actin recycling pathway that is modulated during plasticity (compare [17]).

In conclusion, the here proposed model can serve as a basis to investigate the relation between microscopic properties like molecular dynamics, membrane geometry and emerging properties as spine volume fluctuations. As such it can be extended into various direction.

## Supporting information

**S1 Appendix. Estimating the concentration of available actin**.

**S2 Appendix. Calculating 2D membrane forces**.

**S3 Appendix. Calculating 3D membrane forces**.

## Acknowledgments

We thank the German Science Foundation through the Collaborative Research Center SFB-1286 (Project C3) for funding this research.

